# Tri-n-butyl phosphate inhibits neurogenesis and motor functions during embryonic development in zebrafish

**DOI:** 10.1101/2023.10.27.564317

**Authors:** Gourav Chakraborty, Kedar Ahire, Bhgyashri Joshi, Chinmoy Patra

**Affiliations:** Department of Developmental Biology, Agharkar Research Institute, Pune, Maharashtra, 411004, India; S P Pune University, Pune, Maharashtra, 411007, India

**Keywords:** Tri-n-butyl phosphate, metabolites, zebrafish, neurotoxicity, embryogenesis

## Abstract

Tri-n-butyl phosphate (TBP), an organophosphate ester (OPE), is heavily used as a solvent in synthetic resins, inks, and adhesives and as an antifoaming agent in detergents and emulsion products. Further, it is used as a flame retardant and plasticizer in the fabric and electronics industry. Further, TBP is used to extract uranium, plutonium, and thorium. Thus, widespread uses of TBP in industrialized countries led to the release of TBP and its metabolites, dibutyl phosphate (DBP) and monobutyl phosphate (MBP), in the environment, including air, soil, and water, and get detected in human samples. Accumulating these OPEs over time in humans and other animals may develop toxicological effects. The reports also say OPEs contaminants pass through the mother-fetal transmission route and may affect embryonic development. However, the impact of TBP and its metabolites on vertebrate development has been poorly studied. *Ex-Utero* development, short generation time, high fecundity, optical transparency, and genetic and physiological similarity to humans make the zebrafish a preferred model for studying vertebrate development, organogenesis, small molecule screening, toxicological evaluation, etc. Thus, we aim to explore the toxic effects of TBP, DBP, and MBP on vertebrate organogenesis and overall embryonic development using zebrafish as a model organism. Our study found that TBP inhibits neural growth resulting in decreased spontaneous movements frequency at 22 and 28 hpf without altering the somite and overall embryonic growth in lower doses. In contrast, in lower concentrations, DBP-treated embryos showed normal neuronal and embryonic development and increased spontaneous movement frequency at 22 and 28 hpf. Further, we found that in higher concentrations, TBP is teratogenic, DBP is lethal to the embryos, and MBP did not show detectable toxic effects in the developing embryos. Altogether, we found that TBP inhibits neurogenesis, and its metabolite DBP is neuroexcitatory during embryonic development.

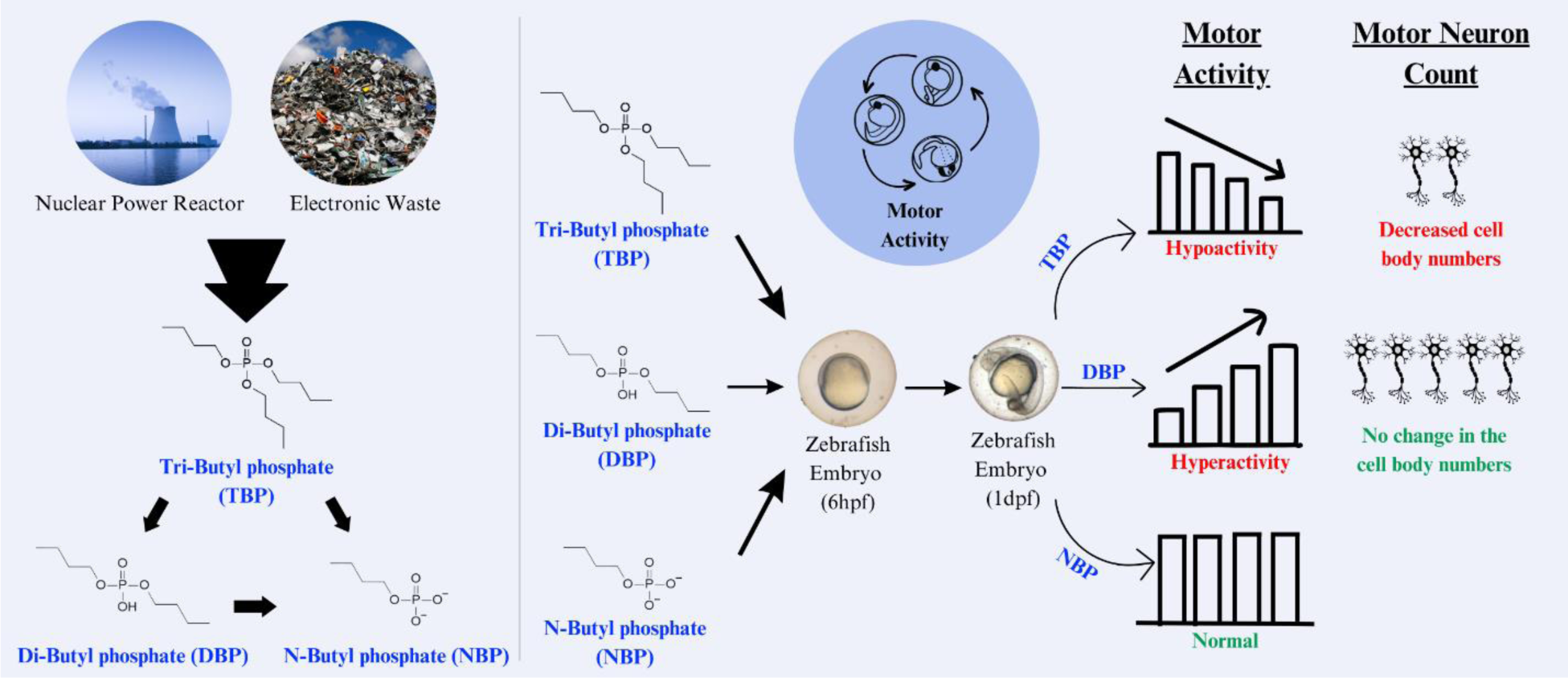

## Introduction

Tri-n-Butyl phosphate (TBP), an organophosphate ester (OPE), is used as an additive or a solvent in various industries, including nuclear, electronic, paper, fabric, plastic, and paint industries. More specifically, TBP is used as a fire-repellant for cellulose fabrics, an antifoaming agent in liquid detergents, emulsions, paints, and adhesives, solvent for cellular esters (van der Veen & de Boer, 2012; Wei et al., 2015), in fire-resistant hydraulic fluids for airplanes, food packaging materials, floor waxes (Larsson et al., 2018; Mendelsohn et al., 2016; van der Veen & de Boer, 2012), etc. Recently the use of TBP has increased exponentially to match the recent demands of fire safety protocols after the ban on the use of polybrominated diphenyl ethers from the markets of the European Union and the United States (van der Veen & de Boer, 2012). Further, in nuclear industries, TBP is used as a solvent to extract uranium, plutonium, and thorium (Kalinowski & Meppen, 2011; WHO 1991, 1991). The global use index of OPEs was 465,000 tons in 2006 (van der Veen & de Boer, 2012), which raised to 680,000 tons in 2015, with an increased usage rate of 7.9% annually (Zeng, Wu, et al., 2018). Reports suggested the presence of OPEs in water (Reemtsma et al., 2008), sediments (Zeng, Hu, et al., 2018), drinking water (Ding et al., 2015), indoor dust (Hoffman et al., 2015), and wastewater treatment plant (WWTP) effluent (Marklund et al., 2005; Sundkvist et al., 2010). The reason behind the presence of OPEs in the air, drinking water, and foods is that they are not covalently bound to indoor and outdoor products (Hou et al., 2021; Wang et al., 2017; Wei et al., 2015), which tend to get released in the surrounding micro-environments due to volatilization, abrasion, and leaching (Hou et al., 2021; Wei et al., 2015).

Further, the radiolysis of TBP happens before complete combustion, i.e., at 700° C, which results in the formation of primary metabolite Dibutyl phosphate (DBP), and subsequent degradation result in the formation of Monobutyl phosphate (MBP) (American Conference of Governmental Industrial Hygienists (ACGIH), 1999; Committee on Updating of Occupational Exposure Limits & Netherlands, 2005; Kalinowski & Meppen, 2011). Open burning of the e-waste using unethical environmental norms also releases TBP and its metabolites as dust in the environment. Similarly, effluents from nuclear power reactors, the paper industries, and e-waste carrying OPEs pollute the nearby water bodies and the environment (Marklund et al., 2005; Sundkvist et al., 2010; Suzuki et al., 1984b). Subsequently, these chemicals are exposed to different organisms living in the water and surrounding via dermal contact, ingestion of dust, inhalation, and food (Bai et al., 2019; Chen et al., 2019; Hou et al., 2020, 2021; Suzuki et al., 1984b). Reports suggest that ingested TBP break into its metabolites DBP and MBP in the liver and kidney of the organisms (Suzuki et al., 1984b), as their presence was detected in urine and blood samples of the organisms (Hou et al., 2020, 2021; Suzuki et al., 1984b). Moreover, several studies showed that TBP and its metabolites diffuse freely through the placenta into the developing fetus (Ding et al., 2016; Hou et al., 2020). A recent pilot study in Guangdong province, China, showed the presence of organophosphate esters (especially DBP and diphenyl phosphate (DPHP)) in amniotic fluid and maternal urine with 13% of mothers had estimated daily intake levels higher than the reference dose, i.e., 2400 ng per kg body weight per day (Bai et al., 2019) raised concerns about the detrimental effects of these chemicals on the fetus’s development.

Several studies on adult animals showed inhibitory effects of TBP on butyrylcholinesterase (BuChE) activity in adult hens (Carrington et al., 1990), the conduction velocity of caudal nerve in adult rats (Laham et al., 1983), and neurobehavior of *Caenorhabditis elegans* (Tang et al., 2022). Recent works showed that TBP has reproductive toxicity in *C. elegans* (Zhang et al., 2021) and 100% mortality in zebrafish embryos upon 64 µM TBP treatment (Tran et al., 2021). However, the effects of TBP and its metabolites on embryonic development and organogenesis remain poorly understood.

*In vitro* development, high fecundity, optical transparency, and genetic and physiological similarity to humans make the zebrafish a suitable model for studying organogenesis, vertebrate development, small molecule screening, and toxicological evaluation (Bambino & Chu, 2017; Menelaou et al., 2008), etc. Hence, we aim to explore the effect of TBP and its metabolites as individual molecules on vertebrate morphogenesis using zebrafish embryos as model organisms. In this work, we used zebrafish embryos in intact chorion during the toxicological analyses to provide natural protection, as the embryos remain in their natural environment. A recent study on zebrafish embryos showed that 64 µM TBP treatment led to 100% lethality. (Tran et al., 2021). Hence, we used lower or sublethal concentrations of TBP, DBP, or MBP to explore their effect on organogenesis. We found that in lower doses, i.e., 10-20 µM concentration, the parent compound TBP inhibit motor neuron growth resulting in decreased spontaneous movement frequency in the pharyngula stage and reduced heart rate at 2 days post fertilization (dpf) without affecting the growth of the embryos. In higher doses, i.e., 40-60 µM in embryo water, TBP was found to be teratogenic but not lethal to the embryos. In contrast, DBP treatment did not show any visible phenotypes, including in neuronal growth until 20 µM concentration; however, DBP-treated embryos showed an increased spontaneous movement frequency. In 40 to 60 µM, DBP was found to be more lethal to the embryos compared to TBP. Notably, MBP-treated embryos until 60 µM remained indistinguishable from their control-treated siblings. Overall, our study on zebrafish found that during early embryonic growth, TBP is a neuronal growth inhibitor, while DBP is likely a neuroexcitatory molecule, and MBP is a non-toxic substance.

## Materials and methods

### Ethics statement

Zebrafish maintenance, experimental care procedures, and protocols employed in the present study were as per guidelines of the Purpose of Control and Supervision of Experiments on Animals (CPCSEA) Committee, Government of India, and approved by the Agharkar Research Institute’s Animal Ethics Committee (IAEC).

### Zebrafish husbandry

The experiments performed were conducted on no background wild-type AB strain (Tran et al., 2021) and *Tg(mnx1:EGFP)* transgenic line, which expresses EGFP in the motor neurons (Nagar et al., n.d.; Saint-Amant, 2006). Embryos ranging from fertilized eggs to 48 hpf were used in this study. The zebrafish’s housing and maintenance were achieved per the pre-recorded protocols (Kimmel et al., 1995; Westerfield, 1993). The study of embryogenesis and the toxicological disturbances in the growth were followed as per the established guidelines (Kimmel, 1989; Kimmel et al., 1995). The fertilized eggs were collected and grown at 28⁰C; a total number of not more than 70 embryos were maintained in 25 ml embryo water in a 10 cm Petri plate.

### Chemical treatment

10 mM stock of Tributyl phosphate (TBP) of >99% purity (240494, Merck), or Dibutyl phosphate (DBP) of >99% purity (68572, Merck), or N-Butyl phosphate (NBP, mixture of mono-N-Butyl phosphate and di-N-Butyl phosphate) of >95% purity (808555, Merck), were prepared by diluting in dimethyl sulfoxide (DMSO). To explore the adverse effects of these chemicals on embryogenesis, 6 hpf (shield stage) zebrafish embryos in chorion were treated with 10, 20, 40, or 60 µM TBP, DBP, or NBP. Control embryos received DMSO as vehicle control. For each treatment, 20 embryos in 10 ml embryo water in a well of a 6-well plate were maintained. Phenotypic evaluation and viability study was performed at 22, 28, and 48 hpf.

### Brightfield imaging

Zebrafish embryos were monitored using a Leica M205 FA stereoscope at different times. To study the overall morphology, control or chemical-treated embryos at 48 hpf were tranquilized in 0.001% Tricaine methanesulfonate (MP Biomedicals), and lateral views of the embryos were captured on a Leica M205 FA stereoscope.

### Spontaneous tail coiling (STC) assay

Spontaneous movement in zebrafish embryos begins at 17 hpf and decreases as the organism matures (Saint-Amant, 2006; Spitzer, 2004). Spontaneous tail coiling frequency (Luo et al., 2022) was assessed to study the embryos’ motor behavior at early development. Three minutes video of each embryo inside the chorion was captured at 22 and 28 hpf at 13 frames/second using the Leica M205 FA stereoscope. Spontaneous tail coiling frequency was counted manually by Fiji. For each condition and time point, 40 embryos, 10 from each independent experiment, were analyzed and plotted the graph using GraphPad Prism 8 software.

### F-acting staining and muscle structure analysis

Whole-mount Alexa 488 conjugated phalloidin (Invitrogen) staining was performed on 28 hpf control and treated embryos to assess the muscle structure. 28 hpf embryos were fixed in 4% paraformaldehyde overnight at 4^°^C, and fixed embryos were washed with PBS several times. PBS-washed embryos were permeabilized with 0.5% Triton X-100, blocked for 1 h in a blocking solution (5% goat serum (MP Biomedicals)/0.5% Triton X-100/ PBS), and incubated with Alexa 488 conjugated phalloidin (1:400) (Invitrogen) for 2 hours at room temperature. Stained embryos were washed with 0.2% Tween 20 in PBS and mounted in 50% glycerol (Merck) on a glass slide. Lateral views of the somatic region dorsal to the yolk sac extension were captured on a Leica SP8 laser scanning microscope, and images were processed and analyzed with LAS X (Leica Microsystems, Germany), keeping all the parameters fixed for scanning the test and control samples. The acquired confocal z-stacks were processed and analyzed with LAS X (Leica) or GIMP (GIMP Development Team) software.

### Confocal microscopy of the motor neuron

Control or chemical-treated wild-type (*mnx1*:EGFP) embryos (Flanagan-Steet et al., 2005) expressing EGFP in the motor neurons were imaged on a laser scanning microscope to explore the motor neuron growth. For confocal microscopy, 28 hpf DMSO (control) or chemical (test) treated *mnx1:EGFP* embryos were embedded in 0.2% low melting agarose (Merck) on a glass bottom Petri plate (Cellvis, USA). Lateral views of the somatic region dorsal to the yolk sac extension of the embedded immobilized live embryos were imaged on a Leica SP8 laser scanning microscope (Leica Microsystems, Germany), keeping all the parameters fixed for scanning the test and control samples. The optical sections of each whole mount organism were merged at maximum projection using Leica LAS AF Lite software (Leica) and processed using GIMP (GIMP Development Team) software.

### Touch-evoked escape response (TEER) assay

To explore the touch responsiveness, 48 hpf of at least 13 control or test-treated embryos were transferred to a 60 mm Petri plate having 10 ml embryo water. All embryos were brought to the center of the plate by creating a centripetal force and allowing them to settle. Tactile stimuli were applied by striking the center of the population with a 10 µl micropipette tip, and their movement was recorded under a Leica M205 FA stereoscope. The percentage of the organisms moving out of the microscopic field view due to the applied tactile stimuli was calculated and plotted the rate in the graph from four independent experiments/treatments using GraphPad Prism.

### Heartbeat frequency analysis

To capture heart functions, 48 hpf control or test molecule treated embryos were placed in a 60 mm petri plate having 10 ml embryo water, and heart movement of each embryo was captured under a Leica M205 FA stereoscope over 10 Sec at room temperature as described earlier (Ma et al., 2021). The heartbeat frequency of 40 embryos, 10 from each independent experiment, was counted manually for each treatment condition and plotted the graph using GraphPad Prism software.

### Statistical analysis

The Student’s T-test was employed to analyze statistical differences in spontaneous tail coiling, touch-evoked escape response, heartbeat frequency, and survival assay. Statistical significance was considered at p<0.05. GraphPad Prism7 software was used to process the data and determine statistical significance. All values are represented as the mean ± s.e.m.

## ACKNOWLEDGMENTS

We are grateful to Prashant K. Dhakephalkar, Director ARI, for his support in performing this work. We thank Satish Bojja for the excellent fish care and ARI for internal support.

## Author contributions

Conceptualization: GC, CP; Methodology: GC, KA, BJ, CP; Validation: GC, BJ; Formal analysis: GC, KA, BJ; Investigation: GC, BJ, CP; Resources: KA, CP; Data curation: GC, BJ, CP; Writing – original draft: GC, CP; Writing – review & editing: GC, CP; Visualization: GC, KA, BJ, CP; Supervision: CP; Funding acquisition: CP

## Results

### Tributyl phosphate inhibits motor activities during embryogenesis

To explore the effect of TBP, zebrafish embryos in the chorion were treated at the blastula stage, the pre-organogenesis stage. For each treatment condition, 20 embryos were maintained in each well of a 6-well plate with 10 ml water, and their morphology and survival rate were observed until 48 hpf (Figure 1 A). Viability analysis showed no lethality in the embryos treated with 60 µM or less TBP (Figure 1 B). Further morphological assessment of the brightfield images of the embryos at 48 hpf found that TBP treatment up to 20 µM does not develop any morphological anomaly (Figure 1 C and S1 A). In contrast, although embryos treated with 40 or 60 µM showed normal somitogenesis, fin fold development, and melanocyte distribution at 28 hpf (Figure S1 B and C), 48 hpf embryos showed a curvature in the caudal region and pericardial edema (Figure S1 A). Next, we wanted to explore the effect of TBP on the motor response of the zebrafish embryos at a concentration that did not show any morphological deformities. Zebrafish start their first motor activity at 17 hpf by the side-to-side mediated movement of their trunk, termed spontaneous tail coiling (STC), which peaks at 19 hpf and gradually decreases with time (Saint-Amant & Drapeau, 1998). These motions result from the activity of the central nervous system during early embryogenesis (Menelaou et al., 2008). Thus, to explore the effect of TBP on central nervous system activity, we captured the spontaneous movement of the test or control-treated zebrafish embryos inside the chorion at 22 hpf and 28 hpf. Quantification of the spontaneous tail coiling over 3 min for each embryo showed significantly decreased movement frequency in 10 or 20 µM TBP-treated embryos compared to their untreated or DMSO-treated siblings at 22 hpf (Sup Video 1-4 and Figure 1 D) as well as at 28 hpf (Sup Video 5-8 and Figure 1 E). The almost abolished spontaneous movement was found in 40 or 60 µM TBP-treated embryos at 22 (Sup Video 9-10 and Figure S1 D) and 28 hpf (Sup Video 11-12 and Sup Figure S1 E). These data indicate that TBP inhibits motor activity at the concentration in which morphological development remains unaffected.

**Figure 1:**
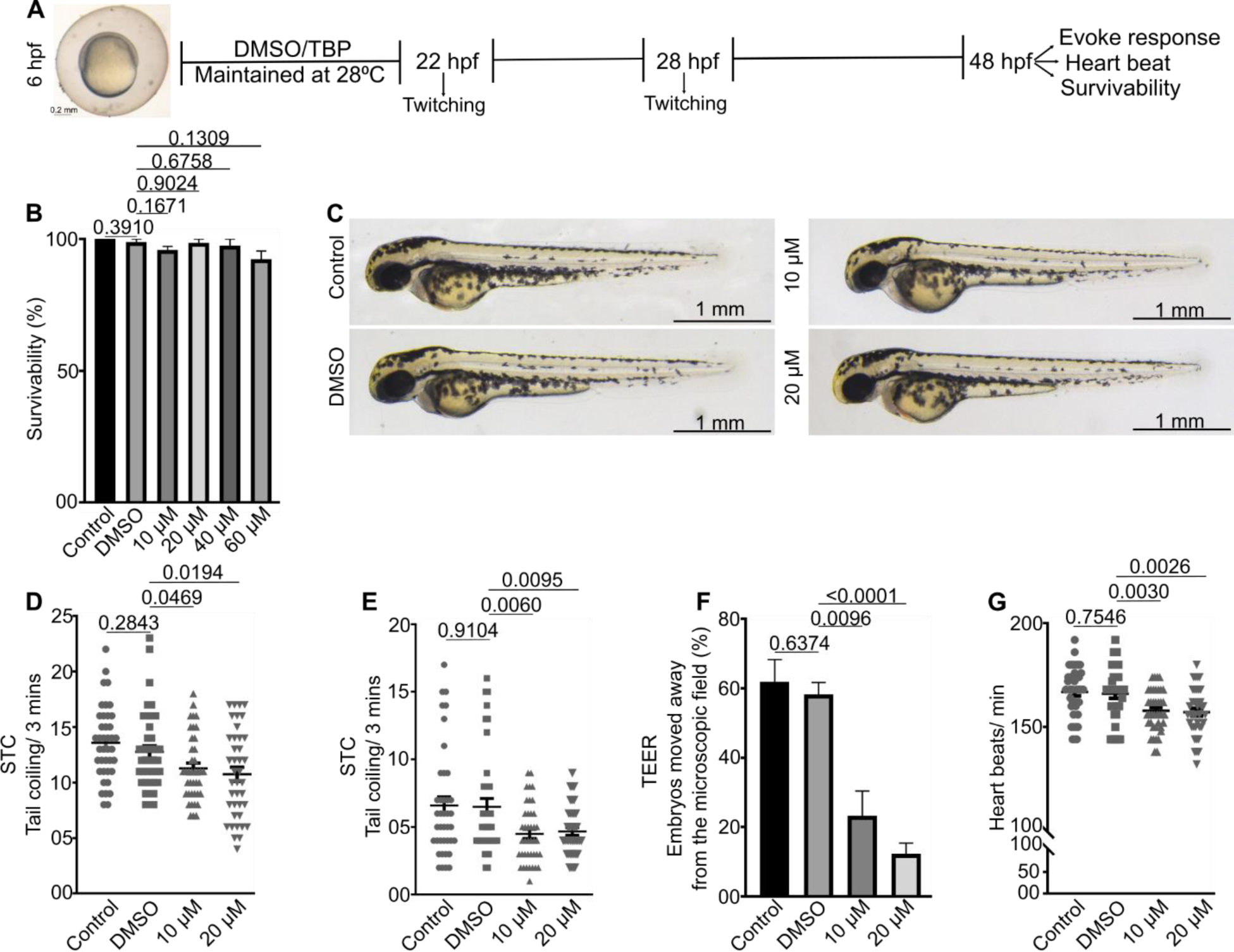
Tri-n-butyl phosphate inhibits motor activities. (A) Schematic representation of the experimentation. (B) Embryo viability at 48 hpf (n=4 sets, 20 embryos in each set). (C) Lateral view brightfield images of the DMSO or TBP treated embryos treated at 48 hpf. (D, E) Dot plots depicting the frequency of spontaneous tail coiling of each organism at 22 hpf (D) and 28 hpf (E) (n=40 from 4 independent experiments). (F) % of embryos moved away from the microscopic field in response to a touch stimulus at 48 hpf (n=4 sets, ≥13 embryos in each set). (G) Heartbeat frequency at 48 hpf (n=40 from 4 independent experiments). Data are mean±s.e.m. in B, D, E, F, and G. The statistical significance of differences was evaluated using a two-tailed Student’s t-test (GraphPad Prism). hpf; hours post fertilization, DMSO; dimethyl sulfoxide, TBP; Tri-n-butyl phosphate.

Upon external touch stimuli, dechorionated embryos start to move contra-laterally at 24 hpf. Around 26 hpf, embryos begin to move forward to a certain extent (Saint-Amant & Drapeau, 1998), and around 48 hpf, they swim freely (Basnet et al., 2019; Saint-Amant & Drapeau, 1998). Since the TBP-treated embryos showed decreased motor activity at 1 day post fertilization (dpf), we sought to explore the effect of TBP on touch response at 48 hpf. Touch evoked escape response (TEER) assay was employed to check the touch receptivity. For each experiment, at least 13 embryos were transferred to a 60 mm Petri plate and swirled to create centripetal force. Once all the embryos reached the center, an external stimulus with the help of a microtip was applied in the middle of the population and captured the animal’s response to the provocation. TEER analysis showed while ∼60% of control-treated embryos moved away from the microscopic field, only ∼25% and ∼15% of the 10 or 20 µM TBP-treated embryos, respectively, moved away from the microscopic field (Sup Video 13-16 and Figure 1 F). Further, none of the ≥ 40 µM TBP-treated embryos moved away from the microscopic field (Sup Video 17-18 and Figure S1 F). Compromised motor activity in TBP-treated embryos persuaded us to explore the effect of TBP on embryonic heart function. Thus, we explored the effect of TBP on heartbeat frequency. 10 and 20 µM TBP has been employed to explore its effect on heart function, as embryos treated with ≤ 20 µM TBP are morphologically indistinguishable from their control-treated siblings. Our analysis at 48 hpf found decreased heartbeat frequency in 10 or 20 µM TBP-treated embryos compared to control embryos (Sup Video 19-22 and Figure 1 G). These data indicate that at higher concentrations, i.e., ≥40 µM TBP is teratogenic and at 10-20 µM TBP negatively influences neuronal activity and cardiac functions without affecting overall growth of the embryos.

### Dibutyl phosphate induces motor activities in zebrafish embryos

Preceding results showed that TBP inhibits motor activities and heart functions. Published evidence indicated ingested TBP break into its metabolites DBP and MBP in the liver and kidney of the organisms (Suzuki et al., 1984a). Thus, we intended to explore the effect of DBP on overall embryonic development and neuronal activity. Zebrafish embryos in the chorion at the blastula stage were treated with DBP or DMSO (vehicle control) and observed their morphology and survival until 48 hpf (Figure 2 A). Viability and morphological analysis showed that treated embryos are viable and morphologically indistinguishable from the control embryos when treated with 10 or 20 µM DBP (Figure 2 B and C). However, ∼25% and ∼75% lethality was observed in embryos treated with 40 or 60 µM DBP, respectively (Figure 2 B). Interestingly, survived embryos remained morphologically indistinguishable from the control embryos (Figure S2), indicating DBP is not teratogenic but lethal to the embryos at ≥ 40 µM.

**Figure 2:**
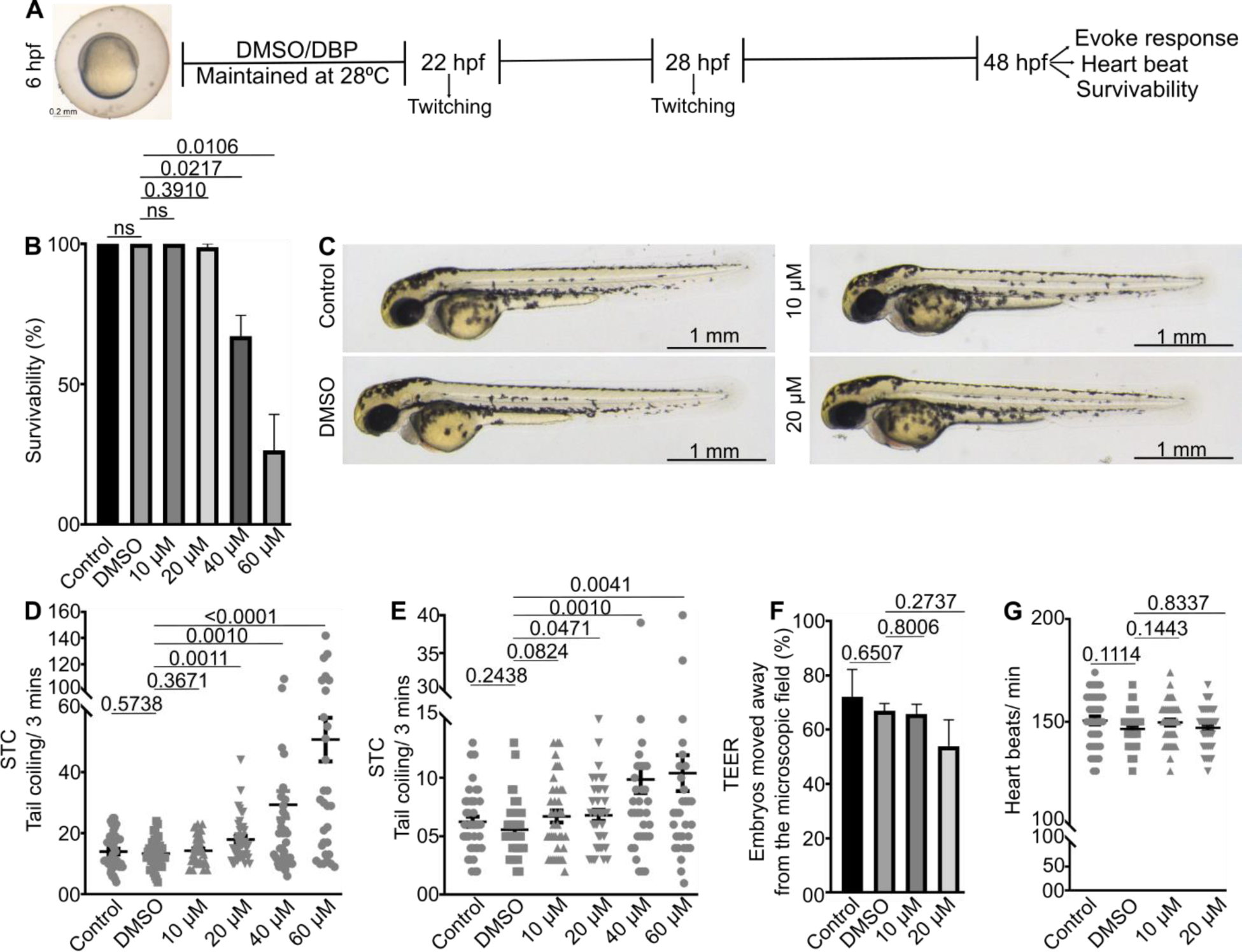
Dibutyl phosphate induces spontaneous tail coiling. (A) Schematic representation of the experimentation. (B) Embryo viability at 48 hpf (n=4 sets, 20 embryos in each set). (C) Lateral views of the DMSO/DBP treated embryos at 48 hpf. (D, E) Dot plots depicting spontaneous tail coiling frequency of each organism at 22 hpf (D) and 28 hpf (E) (n=40 from 4 independent experiments). (F) Quantification of TEER assay indicating % of embryos moved away from the microscopic field in response to a touch stimulus (n=4 sets, ≥13 embryos in each set). (G) Heartbeat frequency at 48 hpf (n=40 from 4 independent experiments). Data are mean±s.e.m. in B, D, E, F, and G. The statistical significance of differences was evaluated using a two-tailed Student’s t-test (GraphPad Prism). hpf; hours post fertilization, DMSO; dimethyl sulfoxide; DBP; dibutyl phosphate.

Next, we sought to find the effect of DBP on the motor responses of the zebrafish embryos by capturing the spontaneous movement of the embryos inside the chorion treated with DMSO (control) or various concentrations of DBP. Spontaneous tail coiling (STC) analysis indicated there is no effect on the STC frequency when treated with 10 µM DBP (Figure 2 D and E). However, in contrast to TBP, DBP-treated embryos showed increased STC in a dose-dependent manner at 22 hpf (Sup Video 23-28, Figure 2 D), as well as at 28 hpf (Sup Video 29-34, Figure 2 E) when treated with 20 µM or higher. Next, we envisioned exploring the effect of DBP on touch stimuli and heartbeat frequency. TEER analysis showed indistinguishable evoke responses to external stimulus among control or 10 µM or 20 µM DBP treated embryos (Sup Video 35-38, Figure 2 F). Further, heartbeat frequency analysis at 48 hpf did not show any effect of DBP on cardiac contraction frequency (Sup Video 39-42, Figure 2 G). These data indicate that, in contrast to TBP, DBP induces spontaneous motor activity in a dose dependent manner and at higher doses lethal to embryos.

### Tributyl phosphate inhibits motor neuron development

Prceeding data showed while TBP-treatment resulting in decreased motor activities, DBP treatment induces motor activities. Thus, we intented to explore the somatic muscle structure in TBP or DBP treated embryos by Alexa-488 conjugated Phalloidin staining on 28 hpf embryos. Microscopical analysis of the phalloidin-stained embryos showed F-actin arrangement in the somite of the 20 µM TBP or DBP-treated embryos are indistinguishable from the DMSO-treated control embryos (Figure 3 A, B), indicating TBP and DBP alter motor activities without affecting somatic muscle morphogenesis.

**Figure 3:**
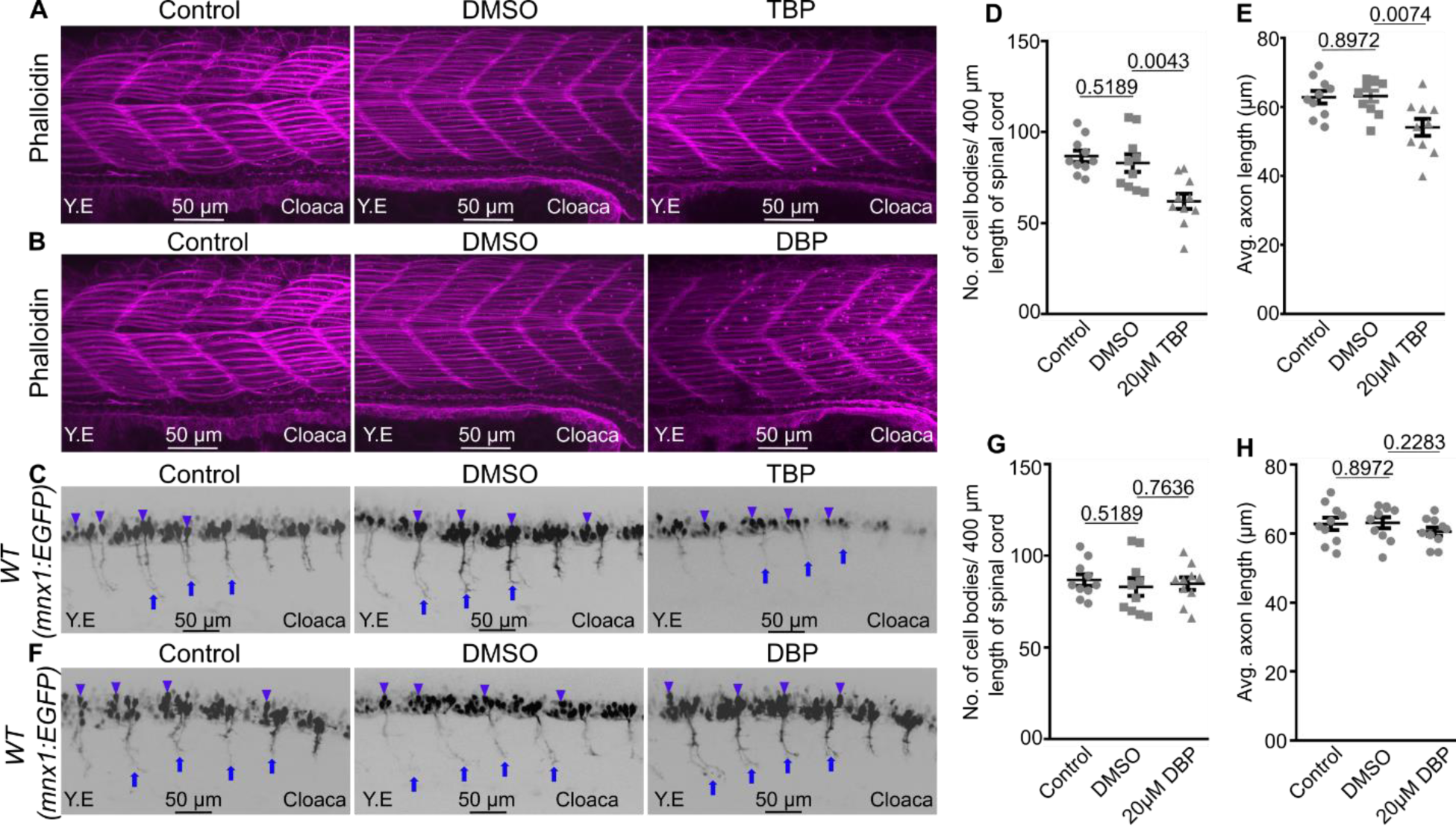
Tri-n-butyl phosphate inhibits motor neuron growth. (A, B) Maximum projections of confocal images of 28 hpf embryos stained for F-actin (Magenta, phalloidin). (C) Maximum intensity projections of confocal images of 28 hpf anesthetized embryos evaluated for motor neurons. Grey indicating motor neurons in the spinal cord. Arrowheads and arrows indicate cell bodies and axonal growth of motor neurons in the spinal cord, respectively. (D, E) Dot plots depicting the quantification of cell body density (D) and axon growth (E) of the spinal cord motor neurons. (F) Maximum intensity projections of confocal images of 28 hpf anasthetized embryos evaluated for motor neurons. Grey indicating motor neurons in the spinal cord. Arrowheads and arrows indicate cell bodies and axonal growth of motor neurons in the spinal cord, respectively. (G, H) Dot plots depicting the quantification of cell body density (G) and axon growth (H) of the spinal cord motor neurons. Data are mean±s.e.m. in D, E, G, and H. The statistical significance was evaluated using a two-tailed Student’s t-test (GraphPad Prism). hpf; hours post fertilization, DMSO; dimethyl sulfoxide, TBP; tributyl phosphate, DBP; dibutyl phosphate.

Development and activation of a primitive spinal circuit is required for the spontaneous contractions of the embryos (Saint-Amant & Drapeau, 1998). Since TBP treatment resulted in a decreasing contractile motion of the animals with an increase in the concentration of TBP (Figure 1 D, E and S1 D, E) without affecting somatic muscle development (Figure 3 A, B), we sought to explore the effect of TBP on motor neuron growth. To determine the motor neuron growth, embryos of Tg(*mnx1*:EGFP) transgenic zebrafish (Flanagan-Steet et al., 2005), (Nagar et al., n.d.), in which EGFP is localized in the motor neurons, facilitating live imaging of motor neuron growth, were used. 6 hpf Tg(*mnx1*:EGFP) embryos were treated with DMSO or 20 µM TBP and imaged the motor neuron development from the spinal cord at 28 hpf. Analysis of the maximum projections of the confocal images showed ∼25% decrease in cell body number (Figures 3 C and D) and ∼15% decrease in axonal growth (Figures 3 C and E) in 20 µM TBP-treated embryos compared to the DMSO-treated control indicating TBP inhibits motor neurogenesis and axonal growth during early embryogenesis.

Next we envisioned to explore the effect of DBP on motor neuron growth as increased STC was observed in the DBP treated embryos in a dose dependent manner (Figure 2 D, E). 6 hpf Tg(*mnx1*:EGFP) embryos were treated with DMSO or 20 µM DBP and imaged the motor neuron development from the spinal cord at 28 hpf. Quantification of the motor neuron showed identical motor neuron cell body density in the spinal cord (Figure 3 F and G) and motor neuron axonal growth from the spinal cord (Figure 3 F and H) between the DMSO-treated control and 20 µM DBP-treated embryos. These data indicate DBP does not have any effect on the motor neurogenesis and axonal growth during early embryogenesis. Altogether, our results showed that during embryogenesis, at ≤ 20 µM, TBP inhibits motor neurone growth and DBP does not have any effect on motor neurogenesis.

### N-butyl phosphate is non-toxic to zebrafish embryos

Preceding analysis on zebrafish embryos revealed TBP and its metabolite DBP have detrimental effects on vertebrate embryogenesis. Hence, we intended to explore the effect of MBP, a metabolite of TBP, on zebrafish embryogenesis and neurogenesis. Since Mono butyl phosphate (MBP) is not available, we have explored the effect of N-Butyl phosphate (NBP, a mixture of mono-N-Butyl and di-N-Butyl phosphate) on zebrafish embryogenesis. Zebrafish embryos in the chorion at the blastula stage were treated with NBP or DMSO (control) and observed their morphology and survival until 48 hpf (Figure 4 A). Viability analysis showed no significant lethality in upto 60 µM NBP treated embryos (Figure 4 B). Further brightfiled images of the embryos at 48 hpf showed upto 60 µM NBP treated embryos are morphologically indistingushible from their control treated siblings (Figure 4 C and S3). These data indicate NBP is not teratogenic or lethal to the embryos.

**Figure 4.**
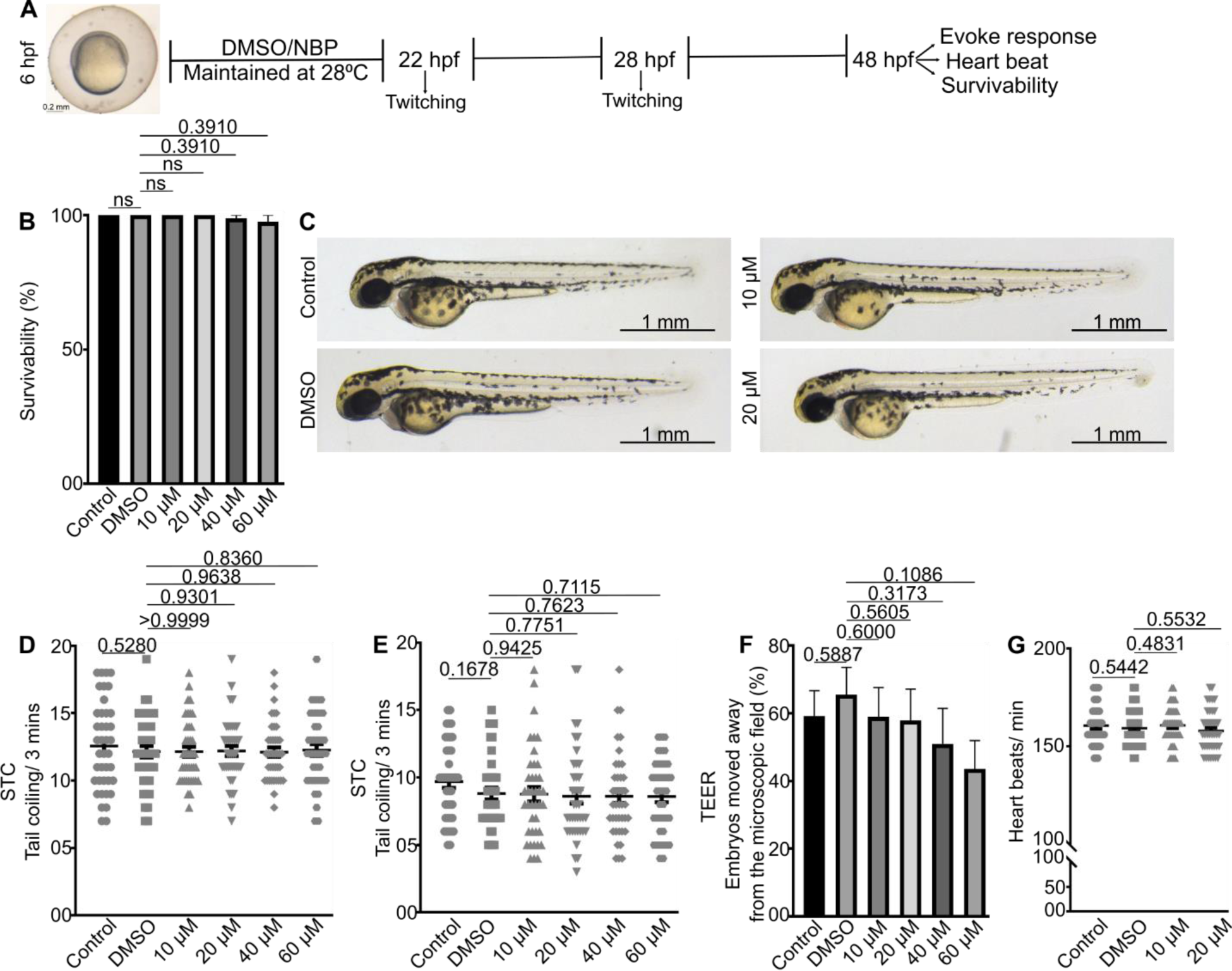
N-butyl phosphate (NBP) is non-toxic to developing embryos. (A) Schematic representation of the experimentation. (B) Bar diagram showing the embryo viability at 48 hpf (n=4 sets, 20 embryos in each set). (C) Brightfield images of DMSO or NBP treated 48 hpf embryos. (D, E) Dot plots depicting the quantification of STC frequency of each organism at 22 hpf (D) and 28 hpf (E) (n=40 from 4 independent experiments). (F) Quantification of TEER assay indicating % of embryos moved away from the microscopic field in response to a touch stimulus (n=4 sets, ≥13 embryos in each set). (F) Heartbeat frequency at 48 hpf (n=40 from 4 independent experiments). Data are mean±s.e.m. in B, D, E, F, and G. The statistical significance of differences was evaluated using a two-tailed Student’s t-test (GraphPad Prism). hpf; hours post fertilization, DMSO; dimethyl sulfoxide; NBP; N-butyl phosphate.

Next, we envisioned exploring the effect of NBP on the motor response of the zebrafish embryo by capturing the STC of the embryos inside the chorion treated with DMSO (control) or various concentrations of NBP. Quantification of the STC frequency revealed NBP does not have any effect on the spontaneous movement frequency at 22 (Sup Video 43-48; Figure 4 D) and 28 hpf (Sup Video 49-54; and Figure 4 E).

Next, we explored the effect of NBP on touch stimuli and heartbeat frequency. TEER analysis showed indistinguishable escape responses to external stimulus among control or upto 60 µM NBP-treated embryos (Sup Video 55-60 and Figure 4 F). Further, heartbeat frequency analysis at 48 hpf did not show any effect of NBP on cardiac contraction frequency (Sup Video 61-64 and Figure 4 G). These data indicate that NBP does not have any significant detrimental effect on neuronal behavior and overall embryogenesis. Altogether, our results showed, in comparison to the TBP and DBP, NBP/MBP is visibly non-toxic to the embryos during embryogenesis.

## Discussion

The experimental outcome of this study on zebrafish embryogenesis identifies organophosphate ester Tri-n-butyl phosphate (TBP) as an inhibitor of motor neuron growth and motor activities. Several observations support this conclusion. First, TBP treatment inhibits spontaneous tail coiling frequency, escape response to the touch stimuli, and heartbeat frequency in lower concentrations, in which the overall development of the embryo visibly remains unaffected. Second, the dose-dependent inhibitory effect of TBP on motor activities and heartbeat frequency. Third, lower concentration TBP treatment showed inhibition of motor neuron growth. In contrast to TBP, DBP-treated embryos showed increased spontaneous movement, however, escape response, and heartbeat frequency remain unaffected. Normal motor neuron growth was visible in both DBP or NBP-treated embryos. In summary, our study on zebrafish embryos finds that TBP is toxic to neuronal growth and teratogenic to developing vertebrate embryos, whereas its metabolite DBP is lethal and NBP is non-toxic.

It has been shown that spontaneous contractions of the embryonic zebrafish musculature are initiated by activation of a primitive spinal circuit, whereas mature locomotive behavior like touch responses is controlled by both the hindbrain inputs and spinal circuits (Saint-Amant & Drapeau, 1998). Rhythmic cardiac contraction and relaxation are regulated by the nervous system from the embryonic stage (Chan et al., 2009). Our observations suggest that TBP treatment reduces spontaneous contraction as well as touch-evoked escape responses. Spontaneous movement, and the locomotive movement of the embryos depends of the embryos, depends on neuronal input and muscle maturation. Our results suggest that TBP treatment does not inhibit muscle growth but inhibits motor neuron growth, which indicates that decreased spontaneous contraction frequency and locomotive behavior in TBP-treated embryos is due to the inhibition of primitive spinal circuit development. Sensory neurons connected to the medulla regulate heart rate. The sympathetic nervous system induces heart rate, whereas the parasympathetic nervous system suppresses heart rate. Our analysis found reduced heartbeat frequency in the TBP-treated embryos.

Organophosphate compounds typically target specific parts of the nervous system, including acetylcholinesterase, nicotine receptors, and muscarinic receptors, which are found in the parasympathetic and sympathetic nervous systems (Menelaou et al., 2008; Robb & Baker, 2023). The parasympathetic effects of organophosphate compounds were found in various systems, including the heart, exocrine glands, and smooth muscles (Dardiotis et al., 2019). Thus, the decreased heartbeat in TBP-treated embryos could be due to the direct effect of TBP on the function of the sympathetic or parasympathetic nervous system, indicating that TBP may have an inhibitory effect on neuronal growth as well as function, which needs to be explored.

Except for the neural toxicity at lower concentrations, at higher concentrations, TBP exposed embryos developed curved body axis in the caudal part, whereas the rest of the body morphologically remained unaffected. Somite formation starts at the end of the gastrulation, continues through the tailbud stage, and is completed by 24 hpf (Stickney et al., 2000). The caudal fin fold initiates to develop from the pectoral fin bud at ∼22 hours post fertilization (hpf) (van Eeden et al., 1996). Since embryos treated with ≥ 40 TBP showed dysmorphic caudal part of the at 2 dpf, indicating that possibly TBP inhibits somite formation or caudal fin fold morphogenesis. However, brightfield images showed that up to 60 µM TBP-treated embryos are identical to their vehicle-treated control siblings at 28 hpf, indicating that TBP does not affect the somite and caudal fin-fold morphogenesis. Defects in axial straightening lead to devastating disorders including scoliosis, characterized by curvatures of the spine (Cheng et al., 2015). Abnormal cerebrospinal fluid (CSF) flow has been implicated in the development of idiopathic scoliosis (Grimes et al., 2016). Thus, it is possible that TBP at higher concentrations inhibits ependymal cell cilia development and cerebrospinal fluid (CSF) flow, resulting in caudal body curvature, which needs to be explored further. In conclusion, our study shows that 10-20 µM TBP inhibits neurogenesis and motor activities and in higher concentration in addition to the neuronal phenotype it likely affects ependymal cell cilia morphogenesis or CSF flow.

In contrast to TBP, it’s metabolite DBP induces spontaneous tail coiling frequency in a dose dependent manner, without affecting the motor neuron growth. Further, DBP treatment did not show any effect on the TEER and heart function of the embryos. Spontaneous contractions of the embryonic zebrafish musculature are initiated by activation of a primitive spinal circuit, whereas mature locomotive behavior like touch responses is controlled by both the hindbrain inputs and spinal circuits (Saint-Amant & Drapeau, 1998). The increased STC in DBP treated embryos having physiological neural circuits indicating that DBP does not have any effect on the neuronal growth but it induces firing rate of developing neurons but not on mature locomotive behavior, which is regulated by the hindbrain and spinal circuits together. Studies showed abnormal firing is implicated in several neurological or mental disorders, including depression and epilepsy (Shao et al., 2021). Thus, embryos exposed to DBP may develop neurological disorders in adult stage without any visible morphological defects.

This study increases our understanding of the detrimental effects of TBP and its metabolites during embryogenesis, which provides their potential hazard to humans that are exposed to these chemicals.

## Conclusion

In summary, this study showcases the effect of alkyl non-halogenated organophosphate ester TBP and its metabolites in vertebrate embryogenesis. We found TBP is neurotoxic and teratogenic. At a lower dose (starting from 10 µM), TBP-treated organisms showed decreased cell bodies and short axonal length of motor neurons along with reduced motor activities. At 48 hpf, we found decreased touch receptivity of the TBP-treated organisms compared to their DMSO-treated control siblings. At higher doses, along with the neurological deformities, TBP-treated embryos showed morphological deformities, including a curved body, and pericardial edema.

In comparison, DBP, the primary metabolite of TBP, showed different toxicological effects on the embryos. DBP-treated embryos showed an increased frequency of spontaneous tail coiling (STC) in a dose-dependent manner. Since the number of cell bodies in the spinal cord and axonal length of the motor neurons in the DBP-treated embryos were identical to their DMSO-treated control embryos, we inferred that the increased STC is likely due to the increased firing rate of developing neurons in DBP exposed embryos.

The secondary metabolite, MBP/NBP, didn’t show any adverse effects on the organisms grown in 10-60 µM. Thus, this study found that while the mother compound TBP inhibits neurogenesis, its primary metabolite DBP induces neural stimuli, and its secondary metabolites MBP/NBP does not have visible effects on the neurogenesis or motor activities of developing embryos, which provides their potential hazard to humans that are exposed to these chemicals.

## Supplementary Information

**Figure S1:**
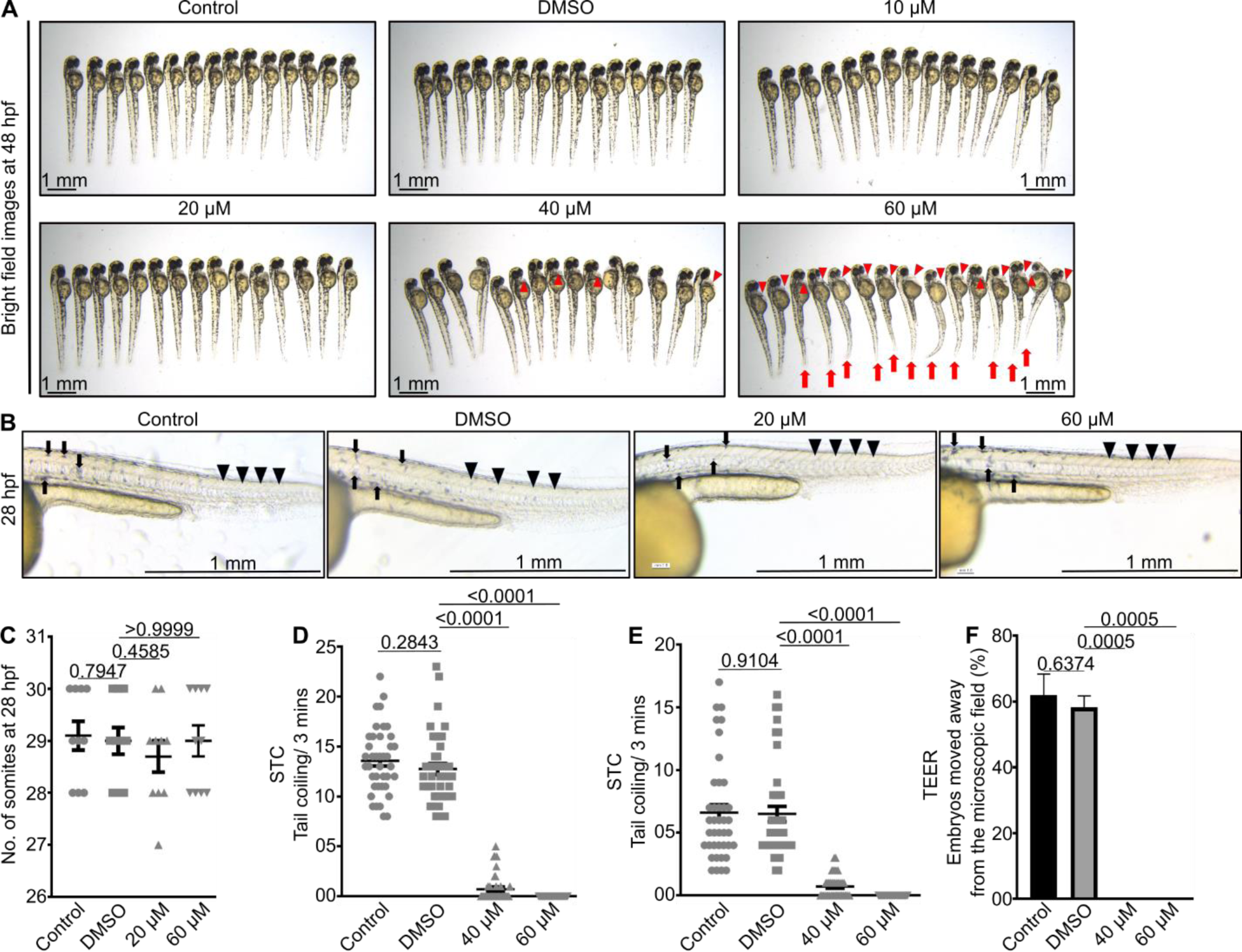
Tri-n-butyl phosphate is teratogenic at ≥40 µM. (A) Lateral views of 48 hpf embryos treated with DMSO or TBP. Red arrows and arrowheads indicate caudal body curvature and pericardial edema, respectively in TBP treated embryos. (B) Representative brightfield images of 28 hpf embryos showing visibly normal growth of the TBP treated embryos. Black arrows and arrowheads indicate melanocytes and somites, respectively. (C) Quantification of somite numbers at 28 hpf. (D, E) Dot plots depicting the STC frequency of each organism at 22 hpf (D) and 28 hpf (E) (n=40 from 4 independent experiments). (F) Quantificatification of TEER assay indicating % of embryos moved away from the microscopic field in response to a touch stimulus (n=4 sets, ≥13 embryos in each set). Data are mean±s.e.m. in C-F. A two-tailed Student’s t-test was employed to evaluate statistical significance of differences (GraphPad Prism). hpf; hours post fertilization, DMSO; dimethyl sulfoxide; TBP; Tri-n-butyl phosphate.

**Figure S2:**
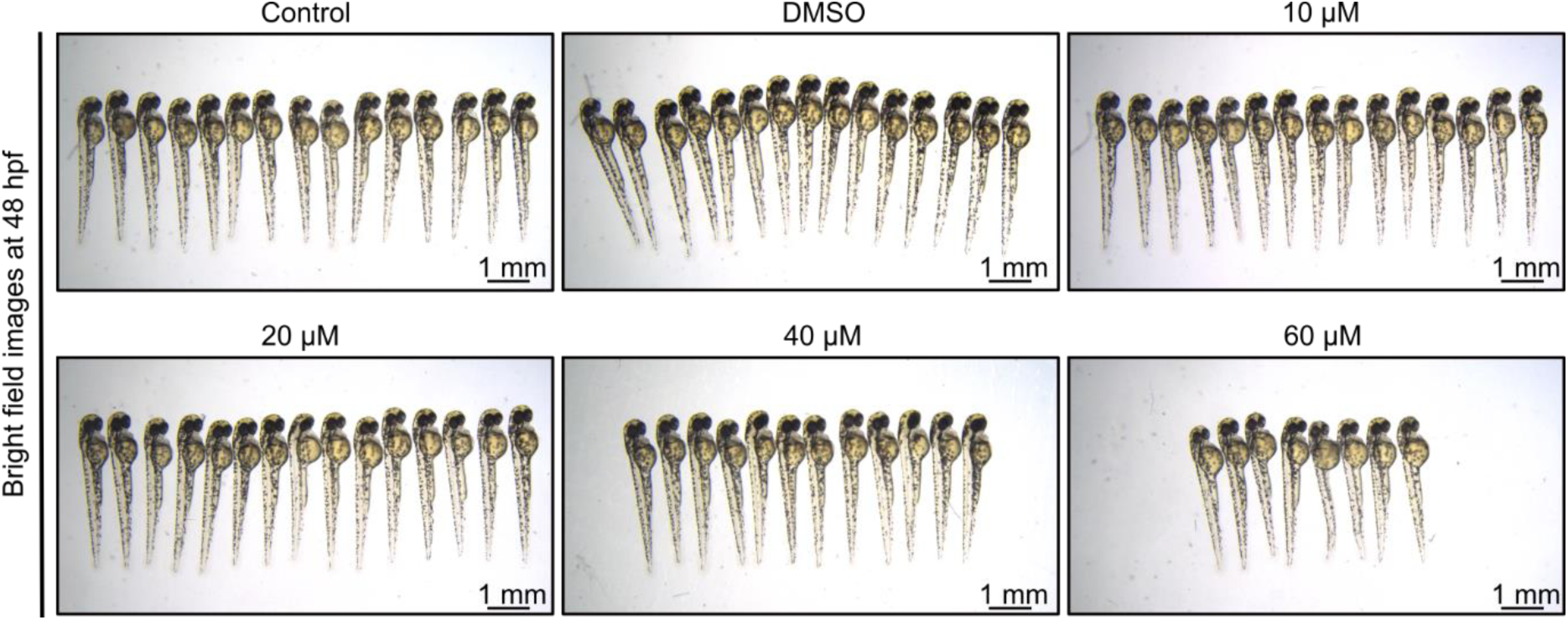
Dibutyl phosphate (DBP) is lethal to embryos at ≥40 µM. Representative brightfield images of 48 hpf embryos.

**Figure S3:**
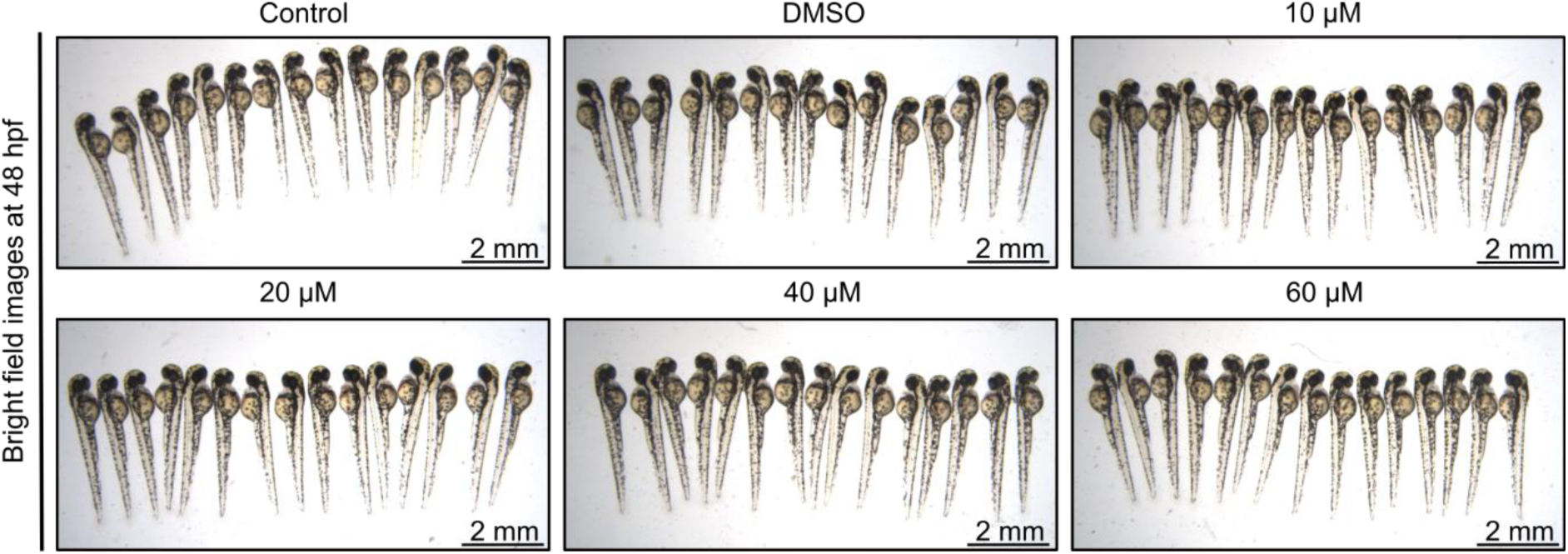
N-Butyl phosphate (NBP) is non-toxic to embryos at ≥40 µM. Representative brightfield images of 48 hpf embryos.

